# Regulation of transcription reactivation dynamics exiting mitosis

**DOI:** 10.1101/2020.04.15.042853

**Authors:** Sergio Sarnataro, Andrea Riba, Nacho Molina

## Abstract

Proliferating cells experience a global reduction of transcription during mitosis, yet their cell identity is maintained and regulatory information is propagated from mother to daughter cells. Mitotic bookmarking by transcription factors has been proposed as a potential mechanism to ensure the reactivation of transcription at the proper set of genes exiting mitosis. Recently, mitotic transcription and waves of transcription reactivation have been observed in synchronized populations of human hepatoma cells. However, the study did not consider that mitotic-arrested cell populations progressively desynchronize leading to measurements of gene expression on a mixture of cells at different internal cell-cycle times. Moreover, it is not well understood yet what is the precise role of mitotic bookmarking on mitotic transcription as well as on the transcription reactivation waves. Ultimately, the core gene regulatory network driving the precise transcription reactivation dynamics remains to be identified. To address these questions, we developed a mathematical model to correct for the progressive desynchronization of cells and estimate gene expression dynamics with respect to a cell-cycle pseudotime. Furthermore, we used a multiple linear regression model to infer transcription factor activity dynamics. Our analysis allows us to characterize waves of transcription factor activities exiting mitosis and identify a core gene regulatory network responsible of the transcription reactivation dynamics. Moreover, we identified more than 60 transcription factors that are highly active during mitosis and represent new candidates of mitotic bookmarking factors which could represent relevant therapeutic targets to control cell proliferation.

## Introduction

Proliferating cells show a global downregulation of transcription during mitosis. This results from the combination of three main processes: 1) nuclear envelope breakdown leading to an increase of the volume that transcription factors (TFs) and the RNA polymerases II (RNAPII) can explore and therefore a decrease of their local concentration around gene promoters; 2) major reorganization of chromatin architecture characterized by chromosome condensation, repositioning of nucleosomes in some regulatory regions, loss of long-range interaction between enhancers and promoters and disassembling of topological associated domains (TADs); and, 3) TF-DNA binding inactivation through postranscriptionally regulated phosphorylation. As a consequence, most TFs and the RNAPII are evicted from mitotic chromosomes and RNA synthesis is drastically reduced [1].

In spite of this global decrease of gene expression during mitosis, proliferating cells are able to maintain their cell identity and propagate regulatory transcriptional programs from mother to daughter cells [2]. Mitotic bookmarking has been proposed as a potential mechanism that could be involved in the transmission of regulatory information during the cell-cycle [3]. Indeed, a significant fraction of TFs are able to remain bound to chromatin during mitosis [4]. These mitotic-bound factors (MFs) show faster interactions with mitotic chromatin than in interphase as reduced residence times have been reported. It is believed that non-specific chromatin or protein-protein interactions between MFs and chromosome coating proteins can explain this fast observed dynamics [4, 5]. However, it has been shown for a handful of MFs, known as bookmarking factors (BFs) [6–9], their ability to interact specifically with at least a fraction of their interphase target sites during mitosis, indicating that chromosomes are not as compacted as previously thought [9]. In fact, chromatin accessibility and nucleosomes landscape during mitosis remain unchanged on bookmarked regions bound by known BFs [11, 12]. This ability of BFs to maintain chromatin structure locally could promote a quick transcription reactivation exiting mitosis.

Transcription dynamics during mitosis and early G1 phase has recently been measured by metabolic labeling of RNA (EU-RNA-Seq) in synchronized population of Human Hepatoma cells HUH7 [10]. Remarkably, this study showed a low but detectable transcription activity during mitosis in up to 8000 genes. Furthermore, transcription reactivation occurred in intense waves exiting mitosis and early G1 phase. However, the study did not take into account that mitotic-arrested cell populations progressively desynchronized once the block was released. As a consequence, RNA measurements are performed on mixture of cells at different internal cell-cycle times. Moreover, it is not understood yet what is the precise role of mitotic bookmarking on mitotic transcription and the transcription reactivation waves. Ultimately, the core gene regulatory network driving the precise transcription reactivation dynamics remains to be identified.

In this paper we developed mathematical models and computational methods to address these open questions. First, in order to correct for the progressive desynchronization of cell populations we assumed that there is a stochastic lag time until a cell can restart the cell-cycle progression again. We characterized the distribution of lag times by analyzing how the observed fraction of mitotic cells evolves over time after the mitotic block is released. This allows us to deconvolve the EU-RNA-Seq data and produce gene expression profiles with respect to a cell-cycle pseudotime and classify the different waves of transcription reactivation in relationship with the cell-cycle progression instead of the experimental time. Moreover, we identified the key TFs determining the transcription reactivation dynamics. To do that, we developed an ISMARA-like model [13] assuming that the expression of genes at a given time point of the cell-cycle progression is a linear combination of the activities of all the TFs that can bind on their promoters. By knowing the deconvolved gene expression and integrating data on transcription factors motif affinities, we calculated the activity of every expressed TF and its role in transcription reactivation exiting mitosis. Indeed, this analysis allows us to divide TFs in groups according to their peak of activity with respect to the cell-cycle pseudotime and identify a core regulatory network of TFs responsible of the observed transcription waves. Interestingly, we do not see a strong correlation between known BFs as FOXA1 and the speed at which their target genes are reactivated. However, we identified around 60 TFs that are highly active during mitosis and represent new candidates of mitotic bookmarking factors.

## Results

### Deconvolution of gene expression data from desynchronized cell populations

In 2017, Palozola et al. published a study based on metabolic labeling of RNA (EU-RNA-Seq) of prometaphase synchronized population of Human Hepatoma cells (HUH7) by arresting cell-cycle progression [10] with nocodazole. EU-RNA-Seq experiments were performed to measure newly synthesized transcripts at 0 minutes, 40 minutes, 80 minutes, 105 minutes, 165 minutes and 300 minutes after mitotic block release as well as for an asynchronous cell population. In this study, the authors highlighted the presence of low levels of transcription during mitosis and the fact that housekeeping genes and not cell-specific genes are activated earlier during the mitotic exit. We reanalyzed the EU-RNA-Seq datasets and characterized the expression dynamics at the gene level. By performing k-means clustering on the gene expression profiles, we identified 5 different clusters, presenting diverse transcription reactivation dynamics over the experimental time. Similarly as the authors reported one of these group showed a peak in the expression at 40 minutes, while others showed a later transcription reactivation (see Fig. 1, panel A).

**Fig 1:**
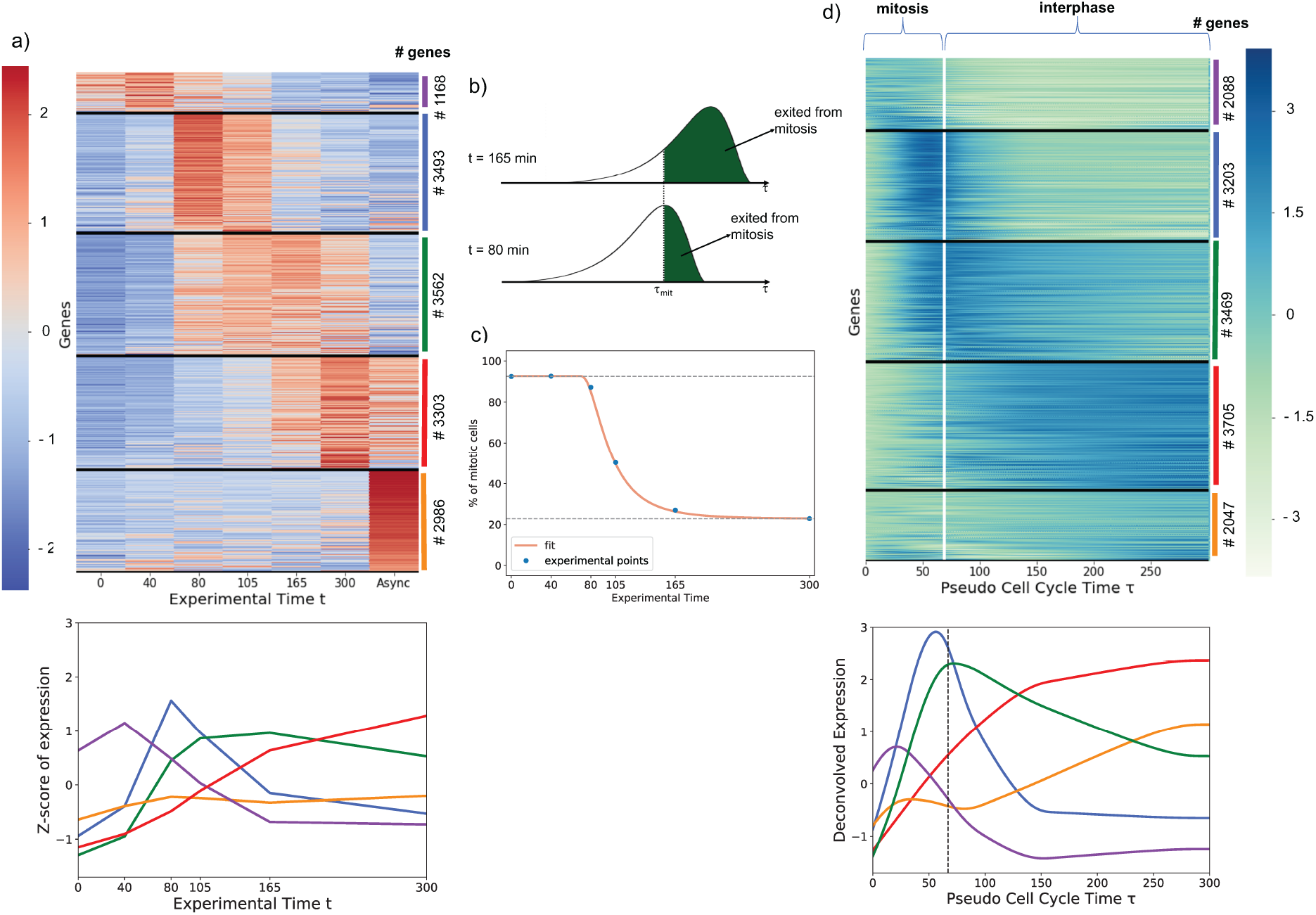
Deconvolution of gene expression data of synchronized cell population leads to dynamic expression profile respect to cell-cycle average profile. **A**: Genes can be divided in groups according to their dynamic expression profiles over time. Each row corresponds to a gene. Black horizontal lines divide the different clusters of genes. The color scale represents the level of the Z-score of expression of each gene, as shown by the colorbar on the left. On the right, the number of genes for each cluster is indicated as well as the color corresponding to the cluster expression average shown on the bottom panel. **B**: We assume that cells have to wait a stochastic, log-normally distributed lag time to start again the cell-cycle progression after the release of the chemical cell-cycle arrest by nocodazole. Here, a pictorial representation of lag time distribution, where the green part of area represents the fraction of cells that already exited mitosis at two different experimental time points. The dashed line indicates the time *τ*_mit_ that cells need to complete mitosis. **C**: Blue dots: quantification by imaging of cells showing condensed (mitotic) and decondensed (non-mitotic) chromatin after mitotic block release (data from [10]); Orange line: model fitting used to infer *τ*_mit_ and the parameters of the log-normal distribution, *σ* and *μ*. **D**: After the deconvolution, genes were divided in groups according the their dynamic expression profile over the cell-cycle pseudotime *τ*. The vertical white line represents *τ*_mit_ and indicates the transition between mitosis to interphase in early G1 phase. The color scale represents the level of the Z-score of expression of each gene, as shown by the colorbar on the right. On the right, the number of genes for every cluster is indicated as well as the color corresponding to the cluster expression average shown on the bottom to one of the curves on the bottom.

Notably, the study did not consider that mitotic-arrested cell populations progressively desynchronize after washing out nocodazole and therefore the reported measurements are performed on mixture of cells at different internal cell-cycle times. In addition, at every experimental time point there is contamination from cells that escape mitotic block. We developed a mathematical model to correct for the desynchronization and the contamination of non-synchronized cells. To do so, we assumed that after mitotic block release there is a stochastic lag time until cells can start again the cell-cycle progression that is log-normal distributed with a certain mean *μ* and standard deviation *σ*. We introduced the concept of *internal cell-cycle pseudotime τ*, defined as the effective cell-cycle time progression of a cell, starting once the lag time is over. We then assumed that there is an average time *τ*_mit_ that cells need to complete mitosis. Finally, we fitted the parameters of the model *τ*_mit_, *μ* and *σ* using data from cell imaging reporting how the fraction of observed mitotic cells evolves over time after the mitotic block is released [10]. This led to an estimated median lag time of 30 minutes and an average time to complete mitosis of 67 minutes (see Fig. 1, panel B and C and Methods for a detailed mathematical derivation).

By applying our model, we deconvolved the time-dependent EU-RNA-Seq data and mapped them onto the internal cell-cycle pseudotime *τ* (see supplementary Table 1). As a result we obtained gene expression dynamics with respect to the cell-cycle progression, allowing us to highlight the transition between mitosis and early G1 phase. Again, we identified 5 different clusters of genes showing distinct transcription reactivation dynamics over the cell-cycle pseudotime *τ*. Strikingly, around 2000 genes showed an expression wave very early during mitosis, presumably around metaphase, while a large fraction of genes reach their reactivation peak just before exiting mitosis, during telophase or during the transition to early G1 phase, as shown in Fig. 1, panel D. In summary, our analysis allows us to correct for desynchronization of cell populations and study gene expression dynamics with respect to the cell-cycle pseudotime highlighting the waves of transcription in relationship with the transition between mitosis and interphase.

### Transcription factor activity dynamics during mitosis and early G1 phase

The transcription waves identified in the previous section are driven by regulatory transcriptional programs mainly activated by transcription factors (TFs). To understand which ones among all TFs are in fact the principal drivers of the transcription reactivation dynamics, we developed an ISMARA-like approach [13]. Thus, we assumed that the normalized log-transformed expression *e_gτ_* of a gene *g* at cell-cycle pseudotime *τ* can be obtained as a linear combination of the cell-cycle dependent activities *A_fτ_* of all TFs *f* that can potentially regulate the gene. The model can be summarized by the following equation:

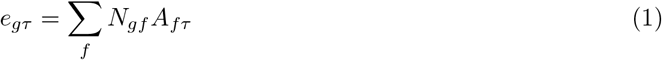

where the values *N_gf_* represent the entries of a matrix **N** containing the number of binding sites for the TF *f* associated with promoter of the gene *g*, taking into account the affinity between the motif of *f* and the sequence of the gene promoter [14]. From the analysis, we excluded TFs associated to unexpressed genes. Furthermore, to avoid overfitting we introduced a regularization term that enforces smooth TF activities over the cell-cycle time and we calibrated using a cross-validation approach. For further mathematical details we refer to Methods.

Our analysis allows us to infer the activity of 332 TFs (see supplementary Table 2). This can be understood as a dimensionality reduction approach as we describe the problem of transcription reactivation with much fewer parameters, since we pass from the analysis of thousands of genes to only hundreds of TFs (see Fig. 2, panel A). To analyze the activity dynamics we use k-means and divide them in 3 clusters, according to their profile over *τ*. We showed that almost 19% of TFs present positive activity during mitosis, with a peak in the first minutes, and then progressive decrease of activity. Conversely, 36% of TFs present a negative activity during mitosis, and then a high activity in early G1. Lastly, the remaining 45% of TFs show a moderate amplitude in their dynamics suggesting that they play a minor role on transcription reactivation dynamics (the results are shown in Fig. 2, panel B). Among TFs that are active during mitosis, we obtain known bookmarking factors as C/EBP, HSF1, TBP, GATA1 and ESRR*β* [3, 10, 15–17] reassuring that our approach is able to identify relevant TFs. Indeed, activates of TFs that are annotated to the Gene Ontology category cell-cycle show an intense dynamics during mitosis and early G1 phase (see Fig. S3). Interestingly, by sorting TFs according to when their highest peak of activity occurs, we observed waves of activity suggesting an intrinsic TF hierarchy with respect to their role on the temporal reactivation of transcription after mitosis (see Fig. 2 panel B).

**Fig 2:**
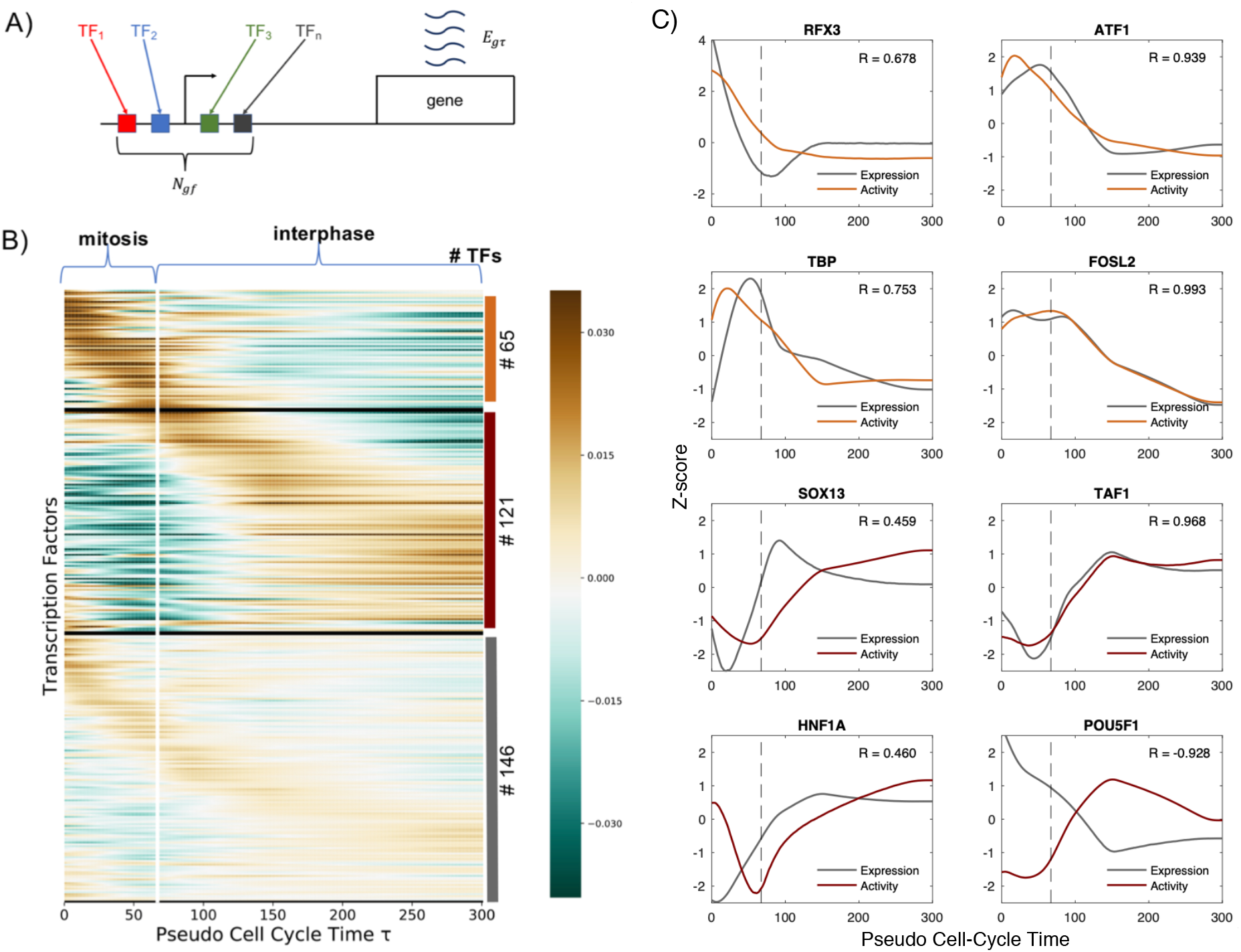
Transcription factor activity dynamics during mitosis and early G1 phase. **A**: Schematic representation of the model: the expression *e_gτ_* of the gene *g* at cell-cycle pseudotime *τ* is a linear combination of the activities of different transcription factors *f* binding the promoter of *g*. *N_gf_* represents the entries of a matrix **N** containing the number of sites for TF *f* associated with promoter of the gene *g*, taking into account the affinity between the motif of *f* and the sequence of the promoter. **B**: TFs can be divided in groups according to their activity dynamics over the cell-cycle pseudotime *τ*. The vertical white line represents *τ*_mit_, and indicates the transition between mitosis and interphase. On the right, the number of TFs belonging to each cluster is indicated. **C**: Activities of mitotic-active (orange curves) and early-G1-active (red curves) TFs that show a high amplitude dynamics. Grey lines show the gene expression dynamics of the corresponding TF genes. Pearson correlation coefficients between the TF activites and expressions are shown in each panel. Dashed lines represent *τ*_mit_.

Determining the molecular mechanisms underlying the TF activity dynamics that we inferred goes beyond the scope of this study. However, we can have a first clue by analyzing the correlation between the TF activity and the expression of the corresponding TF gene. Indeed, a strong correlation indicates that changes in transcription may be responsible for changes in activity. On the contrary, low correlations may suggest that postrasncriptional regulation is required to explain the TF activity dynamics. In Fig. 2 panel C, we show activities of mitotic- and early-G1-active TFs with a high amplitude dynamics together with their expression profiles. Interestingly, TBP, TAF1 and FOSL2 show a high positive correlation indicating that their activities may be regulated at the transcriptional level. On the other hand, SOX13 and HNF1A show a clear delay between expression and activity which could reflect the delay on the accumulation of active protein due to mRNA and protein half-lives or postranscriptional regulation. Strikingly, POU5F1 shows a strong negative correlation which suggests that may act mainly as a repressor. In summary, our analysis not only allows us to identify the activity dynamics of key TFs involved in transcription reactivation but also provides preliminary hints on the molecular mechanisms that may be involved in such dynamics.

### Bookmarking and transcription reactivation kinetic

Next, we investigated the role of mitotic bookmarking in the transcription reactivation dynamics. To do that, we analyzed the expression of genes associated to FOXA1, a liver-specific factor and one of the first identified bookmarking factors. We used mitotic ChIP-Seq data from a study of Caravaca et al. [6]. We selected the genes associated to FOXA1 ChIP-Seq peaks (see Methods) and we calculated the average expression of these genes and compared it with the overall average gene expression. Surprisingly, genes associated to FOXA1 reach their activation peak later than the overall peak of gene expression that occurs during the transition between mitosis and early G1 phase (see Fig. 3, panel A). Then, we compared the activity of FOXA1 with the average activity of all TFs, revealing a negative peak during mitosis (see Fig. 3, panel B), in accordance with the results shown in Fig. 3, panel A. These results suggest that FOXA1, despite its presence on mitotic chromosomes through specific and non-specific interactions, is not sufficient to promote quick transcription reactivation. However it may play a structural function by keeping the chromatin open to promote binding of other TFs.

**Fig 3:**
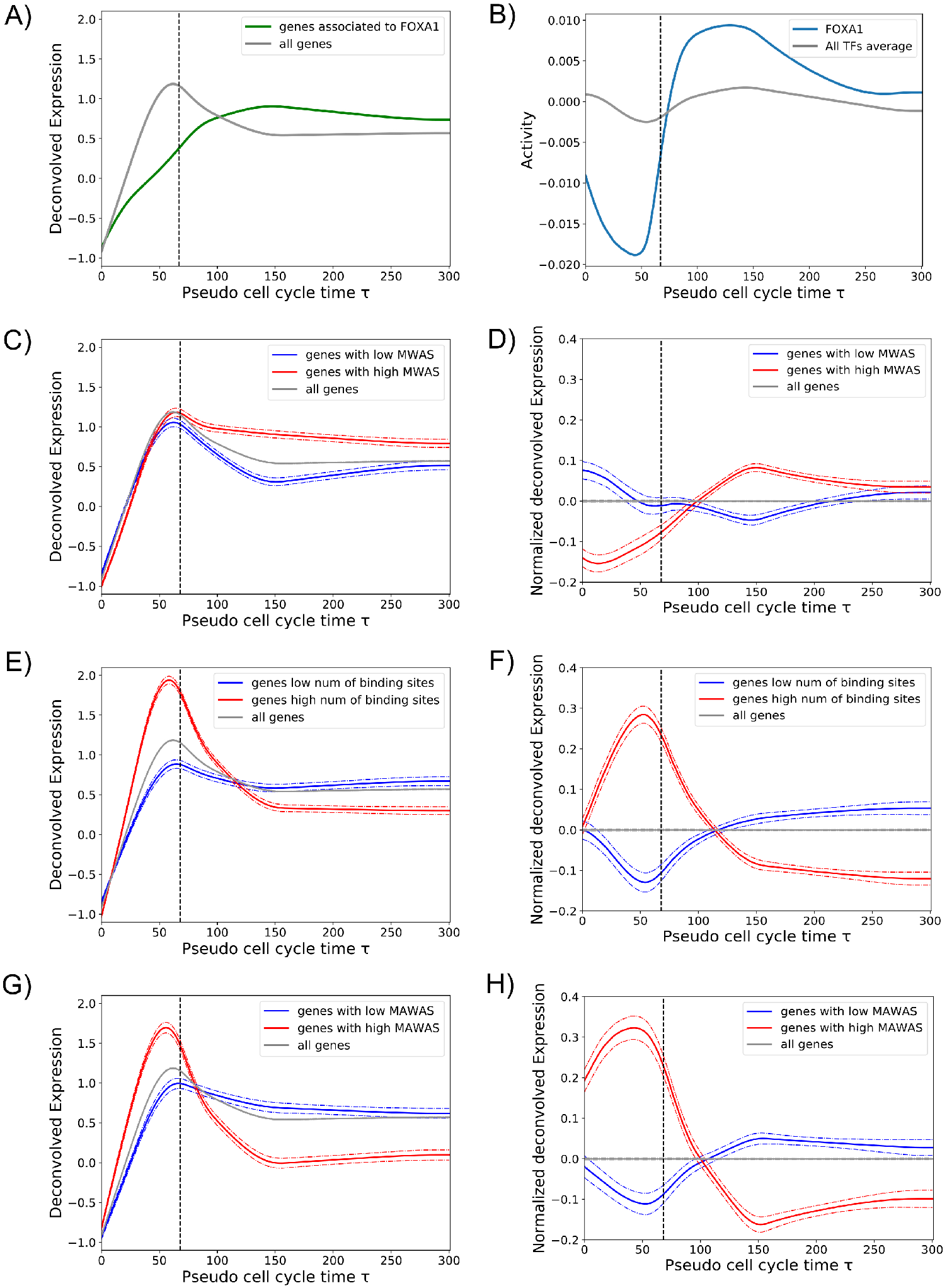
Bookmarking and transcription reactivation dynamics. **A**: The average expression of all genes (grey line) was compared with the average expression of FOXA1 target genes during mitosis (green line). **B**: The activity of FOXA1 (blue line) in comparison with the average activity of all TFs (grey line). **C**: Gene expression pattern as a function of the promoter MWAS (MBF weighted average score). The average expression of all genes (grey line) was compared to the average expression of genes whose promoters tend to be regulated by TFs with high MBF (red line) and low MBF (blue line) (see Methods). **D**: The same as in panel C, but gene expression patterns are shown normalized by the overall average expression (grey line). **E**: Gene expression pattern as a function of the total number of TF binding sites in the promoter. The average expression of genes with large (red) and low (blue) number of promoter binding sites are compared to the overall averege expression (grey line). **F**: The same as in panel E, but gene expression patterns are shown normalized by the overall average expression (grey line). **G**: Gene expression pattern as a function of the promoter MAWAS (mitotic activity weighted average score, xdd Methods). The average expression of genes with high (red) and low (blue) MAWAS are compared to the overall averege expression (grey line). **H**: The same as in panel G, but gene expression patterns are shown normalized by the overall average expression (grey line). Dashed vertical lines in all panels indicate *τ*_mit_.

To scale up this analysis we took advantage of a recent large scale study by Raccaud et al. in 2019. The authors were able to systematically measure the mitotic chromosome binding of 501 TFs in mouse fibroblast cells by live-imaging cell lines carrying exogenous florescence constructs. The mitotic bound fraction (MBF) was defined as the fraction of fluorescence signal located on mitotic chromosomes over the total cell signal. According to this score the TFs were divided in three categories (enriched, intermediate and depleted) indicating their capacity to bind mitotic chromosomes and their potential to be bookmarking factors. We then assumed that human TFs in HUH7 cells behave similar as their mouse paralogs and assigned the corresponding MBF score. We hypothesized that genes regulated by TFs with high MBF should be ready to be reactivated earlier. To test this, we calculated a MBF weighted average score (MWAS) for each promoter as the average MBF of all the TFs that regulate a given gene promoter weighted by the number of their binding sites. Then, we divided genes in high and low MWAS and we calculated the average expression of these two groups (see Methods for further information). Genes associated to high MWAS, i.e gene that tend to be regulated by TFs with high MBF, did not show a faster reactivation dynamics but a significant larger expression during early G1 phase (see Fig. 3, panel C and D). Consistently, we showed no significant difference between the MBF distribution of TFs with high activity during mitosis or during early G1 (supplementary fig. Fig. S4). These results indicate that there is an absence of correlation between TF mitotic binding and TF mitotic activity and quick transcription reactivation of their target genes. We cannot rule out that the absence of correlation could be due to the fact that MBF scores were measured with an artificial system in a different cell line of a different organism. However, the absence of correlation may suggests that that previously reported mitotic bound factors, as in the case of FOXA1, may have a structural function by keeping chromatin open during mitosis and the speed of transcription reactivation may be then regulated by other determinants.

Next, we studied whether promoter architecture could be one of the determinants of early transcription reactivation. Surprisingly, just the total number of binding sites within the gene promoter is a strong feature to predict early or late reactivation. Indeed, average expression of genes with large number of binding sites shows a quick transcription reactivation during mitosis, in contrast to a reactivation during early G1 of genes with small number of binding sites (see Fig. 3, panel E and F). Two non-exclusive mechanisms could explain why strong promoters reactivate earlier: first, gene promoters with more TF binding sites may be easier to kept accessible during mitosis as more TFs could compete against nucleosomes leading to nucleosome free regions. Second, large number of binding sites may facilitates that TFs find the promoters and thus increase the chance to recruit the transcriptional machinery. Finally, as expected, genes that have a large number of binding sites for TFs with a high inferred activity during mitosis (high mitotic activity weighted average score, MAWAS), showed a high mitotic transcription and a quick transcription reactivation (see Fig. 3, panel G and H, and supplementary table 3). Therefore, we believe that our method allows to identify new bookmarking factors that should not only bind mitotic chromosomes but be able to bind specific DNA binding sites during mitosis. In addition, we predict that promoters with large number of binding sites for these TFs should show a higher degree of chromatin accessibility during mitosis.

### Identification of the Core Regulatory Network responsible for the transcription reactivation after mitotic exit

Next, we wanted to identify the TFs, among the 332 for which we are able to infer activities, that have a major role on the reactivation of transcription exiting mitosis. Namely, the key TFs that if perturbed may affect more significantly the measured gene expression patterns. To do so, we calculated the fraction of explained variance as a measure of the performance of our model to fit the data. Then, we defined a TF importance score as the reduction on fraction of explained variance when the TFs is removed as an explanatory variable from the multiple linear regression model (see Methods for further details). A list of all TFs sorted according to their importance score is given in the supplementary table 4.

Furthermore, in Fig. 4 we show a Core Regulatory Network (CRN) where the nodes are formed by the 5% top most important TFs and the links represent potential regulatory interaction between the selected TFs according to the presence of TF binding sites in their promoters. Interestingly, the CRN shows a large number of regulatory links (149 connections) while random networks with the same number of TFs produce a smaller number of connections (36 on average). Thus, the high interconnectiviy of our CRN suggests that the identified TFs may be related functionally. Indeed, some of them have been reported to be involved in cell-cycle or cell proliferation and growth as ATF1, FOS, CEBPZ, SP3 and KLF4 [18]. Moreover, the CRN structure shows multiple feedback loops rather than a hierarchical network as one could expect taking into account the observed sequential waves of transcription reactivation. This type of structure has the potential to show cycling dynamics which may be important not only for the reactivation after mitosis but for the regulation of transcription across the whole cell-cycle. In conclusion, we predict that these TFs could represent relevant therapeutic targets to control cell proliferation.

**Fig 4:**
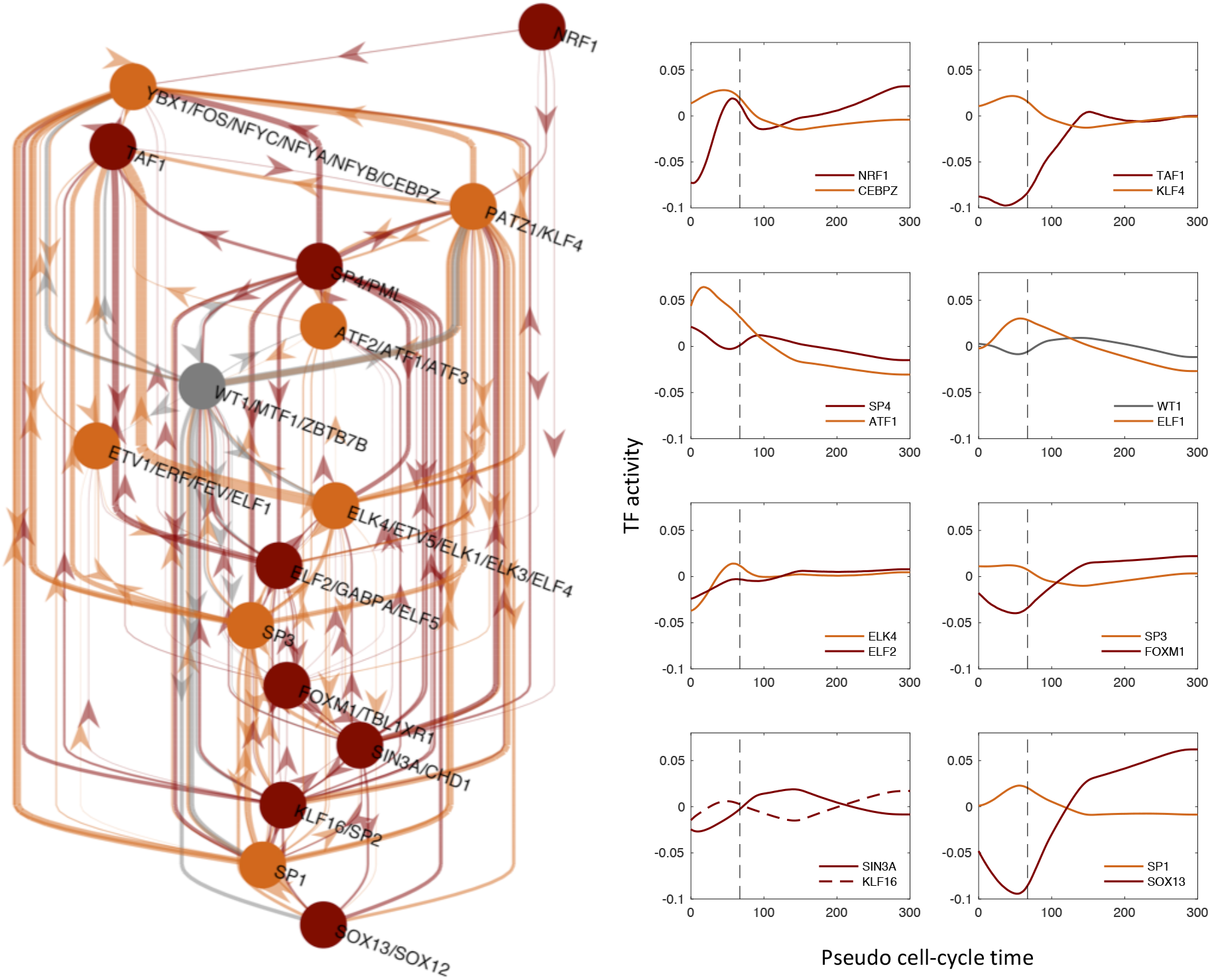
Identification of the Core Regulatory Network responsible for the transcription reactivation after mitotic exit. Inferred core regulatory network (CRN) by selecting the top 5% of the TFs according to their importance in explaining the gene expression patterns and their inferred time-dependent activities. Colours indicate the cluster to which TFs belong, in accordance with the in Fig. 2, panel B. Note that the majority of TF motifs are associated uniquely to single TF, while some other motifs are shared by more than one TF, precluding the inference of single TF activities of motifs which can be potentially bound by more than one TFs.

### Genes within in the same TAD show similar reactivation dynamics

The connection between transcription and the 3D structure of chromatin is currently a very active field of research. During mitosis, topologically associated domains (TADs) are disrupted and rebuilt at different dynamics during the transition between mitosis and G1 phase [20].However, the causal connection between transcription reactivation and chromatin structure reformation exiting mitosis is not known yet. As a first attempt to investigate this relationship, we analyzed the correlation between transcription reactivation profiles of genes belonging to the same TAD. To do that, we took two TAD lists identified in human IMR90 and ES cells, from Hi-C experiments [21]. Although these are different cells lines than the HUH7, it has been shown that TADs are highly conserved between different cell-types and even different organisms [21]. We obtained a total of 2290 and 3061 TADs respectively. For each TAD in both lists, we identified the expressed genes that are located within its limits, finding respectively 10225 and 9849 genes. Pearson correlation coefficients were calculated between all pairs of genes within the same TAD. As a control, expressed genes were randomly located into TADs respecting the total number of genes in each TAD. Distribution of correlation coefficients as well as distribution of random coefficients are shown in Fig. 5. Interestingly, genes belonging to the same TAD show a higher correlation than random pair of genes, indicating that transcription reactivation dynamics are similar between genes that are located near in space within the same local self-interacting chromatin 3D structure. Finally, we hypothesized that TADs containing genes characterized by a quick transcription reactivation should show a faster reformation exiting mitosis. Further experimental work would be required to validate our hypothesis.

**Fig 5:**
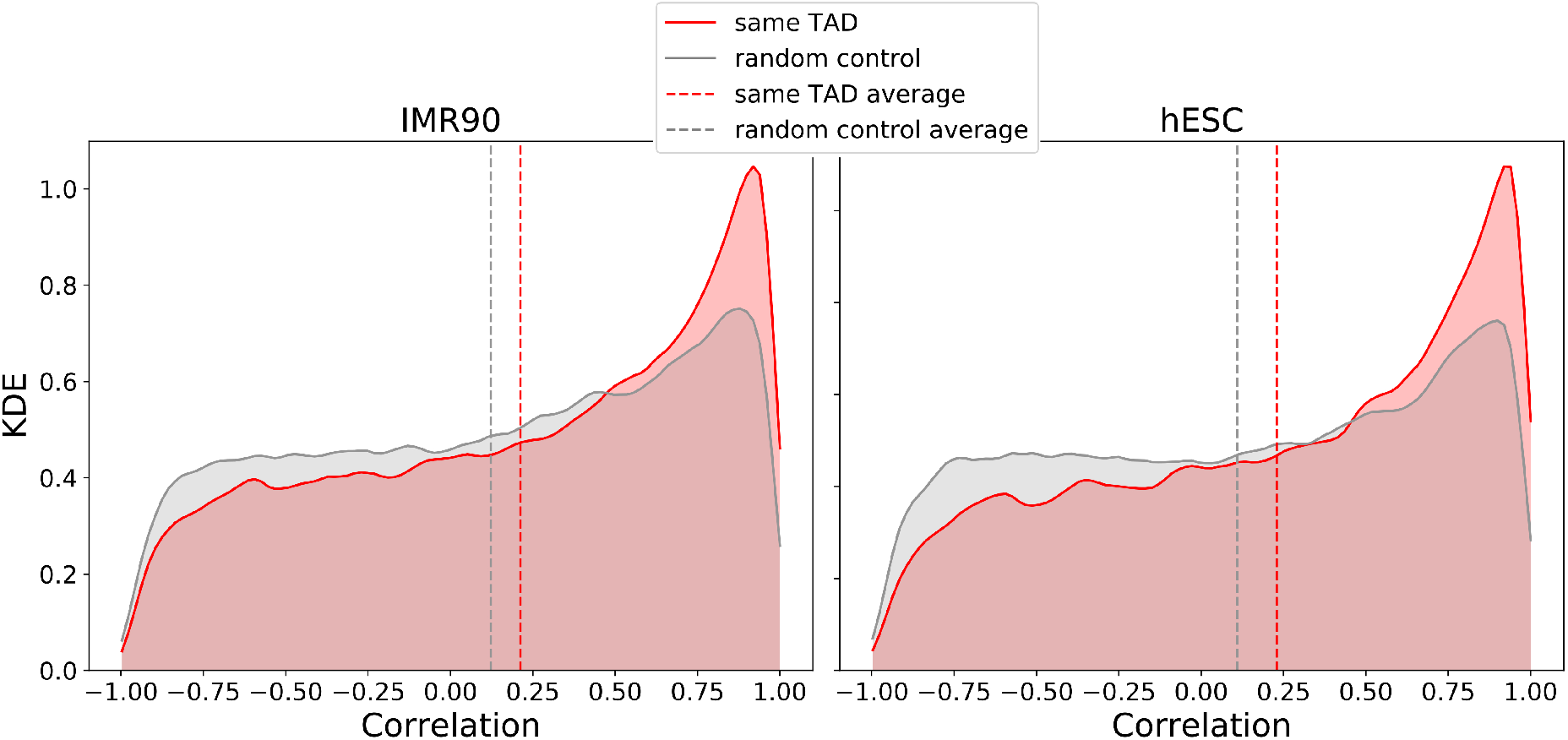
Genes belonging to the same TADs show an higher correlation in expression. Correlations between gene expressions for genes belonging to the same TAD were calculated, for two different TADs sets (IMR90, hESC cells). In red, the KDE of the distribution of the correlation is plotted, in comparison with a random model (grey plot). Dashed vertical lines represent the average values of the corresponding distributions.

## Discussion

Cell identity maintenance in proliferating cells is a biological process that has crucial implications in developmental biology, regenerating medicine and cancer. Nevertheless, the precise molecular mechanisms responsible for the transmission of regulatory information from mother to daughter cells are not fully understood yet. Mitotic bookmarking by TFs through specific DNA binding on mitotic chromosomes has been proposed as a mechanism to reinforce cell identity maintenance during cell division. In this paper we studied the regulation of transcription reactivation exiting mitosis and the connection with mitotic bookmarking.

First, we reanalyzed time-dependent EU-RNA-Seq data on synchronized cell populations by a mitotic arrest, to correct for the progressive desynchronization of cells after block release. This allowed us to estimate gene expression profiles with respect to a cell cycle pseudotime with an explicitly defined transition between mitosis and early G1 phase. Remarkably, we identified a set of genes that show a very early wave of transcription reactivation during mitosis. However, the majority of genes showed a peak of transcription at telophase or during the transition between mitosis and G1.

Next, we estimated TF activity dynamics of 332 expressed TFs by fitting a multiple linear model to the deconvolved gene expression profiles. We observed time-dependent waves of TF activities suggesting an intrinsic TF hierarchy with respect to their role on transcription reactivation after mitosis. In addition, we investigated whether TFs previously reported to bind mitotic chromosomes were responsible for a faster reactivation dynamics. Surprisingly, we did not find a strong correlation between genes regulated by mitotic bound TFs and the speed of reactivation. However, our approach allowed us to identify around 60 TFs that are highly active during mitosis and represent new candidates of mitotic bookmarking factors. Therefore, we predict that the interactions of these factors with their specific target sites during mitosis are the molecular mechanisms responsible for mitotic transcription and transcription reactivation. Moreover, we hypothesize that these specific interactions may also play an important role maintaining chromatin accessibility on mitotic chromosomes. Further experimental work would be needed to validate our hypothesis and predictions.

Moreover, we reconstructed a core regulatory network underlying the dynamics of transcription reactivation exiting mitosis, by selecting the key TFs that showed the highest explanatory power in our multiple linear regression model. Then, we propose a list of candidates to be the crucial players in the process of reactivating the gene expression in the first stages of the interphase, ensuring the cell identity. We predict that these TFs could represent relevant therapeutic targets to control cell proliferation. Further experiments are required to validate our predictions and prove the active role of TFs on chromatin accessibility and 3D structure.

## Methods

### Fitting of model parameters for deconvolution of gene expression data

To estimate gene expression dynamics with respect to an internal cell-cycle pseudotime, we assumed that after the release of the synchronization there is a stochastic lag time until cells can start again the cell-cycle progression. According to our model, the cell-cycle progression of a cell is represented by the internal cell-cycle pseudotime *τ* = *t − η*, where *t* is the experimental time and *η* is the stochastic lag time that the cell had to wait until the cell-cycle progression was restarted again. We further assumed that the lag time *η* is log-normally distributed with a certain mean *μ* and standard deviation *σ*. Then, the probability of finding a cell in the population with an internal cell-cycle pseudotime *τ* at a given experimental time *t* can be written as:

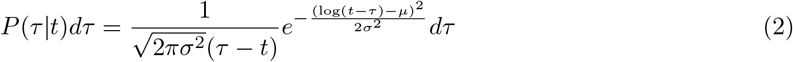

Assuming that cells require an average time *τ*_mit_ to complete mitosis we can calculate the fraction of cells waiting for mitosis to be finished as 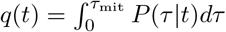 and solving the integral we obtain:

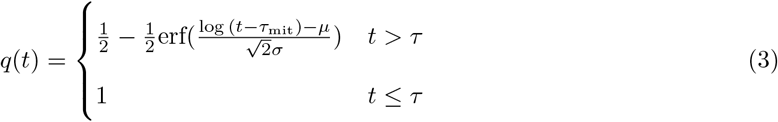

Then, we fit the parameters of the stochastic model (*τ*_mit_, *μ* and *σ*) by using data from [10] on the time evolution of the number of mitotic cells observed after synchronization treatment release. To do so, we define the likelihood of the data based on the assumption that the cell counts follow a binomial distribution with probability *q*(*t*). Thus,

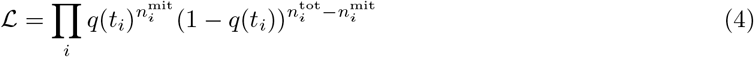

where 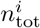 and 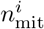 are, respectively, the total number of cells and the number of cells in mitosis counted at experimental time *t*_0_ = 0 minutes, *t*_1_ = 40 minutes, *t*_2_ = 80 minutes, *t*_3_ = 105 minutes, *t*_4_ = 165 minutes and *t*_5_ = 300 minutes. Then, the log likelihood can be written as:

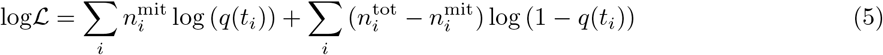

By performing an optimization of Eq. 5 (python *scipy optimize* package, *nelder-mead* algorithm), we can infer the parameters *τ*_mit_, *μ* and *σ*, obtaining, respectively, 67min, 3.43 and 0.74. Once the parameters have been inferred, the probability *P*(*τ*|*t*) is fully determined and, therefore, we can recover the gene expression with respect to the internal cell-cycle pseudotime *τ* using the following convolution equation:

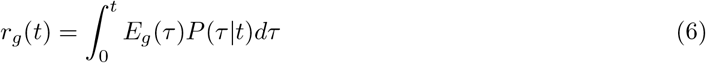

where *r_g_*(*t*) represents the expression of the gene *g* at experimental time *t* (given by the EU-RNA-Seq data), *E_g_*(*τ*) is the expression of the same gene *g* at the cell-cycle pseudotime *τ*. This equation basically reflects that the gene expression measured at a certain experimental time is the population average over the expressions of cells at different cell-cycle times. In case of perfect synchronization over time, the probability *P*(*τ*|*t*) would become a Dirac delta function and the gene expression in both times would be the same. Furthermore, we took into account the fact that the samples were contaminated by a fraction *π_M_* = 0.23 of cells that never exited mitosis and a fraction *π_I_* = 0.075 of cells that did not response to the mitotic block and stayed in interphase (see Fig. 1, panel C). It means that only a fraction *π_C_* = 1 − *π_M_* − *π_I_* starts again the cell cycle progression within the duration of the experiment. Then, this can be summarized by describing the measured gene expression as a mixture of the three cell populations as follows:

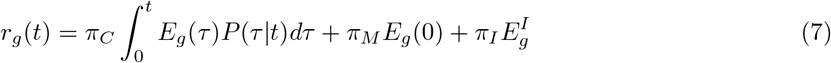

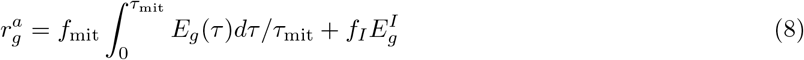

where, 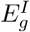 is the average expression during interphase and an extra equation is included to relate the gene expression 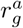 measured on an asynchronous cell population as a weighted average of the gene expression during mitosis and during interphase where the weights reflect the fraction of the cell-cycle duration *T_C_* that cells expend on average in each phase, i.e *f*_mit_ = *τ*_mit_/*T_C_* and *f_I_* = 1 − *f*_mit_.

Then, to perform the deconvolution we discretized the cell-cycle pseudotime into small intervals (*δτ* = 1 min) and expressed the Eq. 7 and 8 into matricial from: **r**_*g*_ = *M* **E**_*g*_, where the expression vectors are defined as 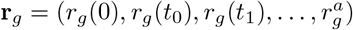 and 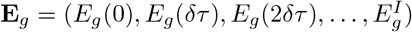 and the matrix *M* is the sum of three components: *M* = *M_C_* + *M_M_* + *M_I_* that account for the three distinct cell populations. First, the cell-cycle matrix *M_C_* models the desynchronization of the cells that re-enter the cell cycle, as in Eq. 6, and can be written as:

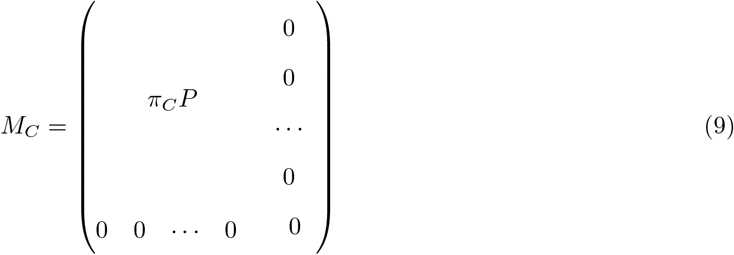

where *P* is the discrete version of Eq. 3. Second, the mitotic matrix *M_M_* adds to the model the contribution of the cells that are still in mitosis by mapping them into *τ* = 0. Its explicit form is:

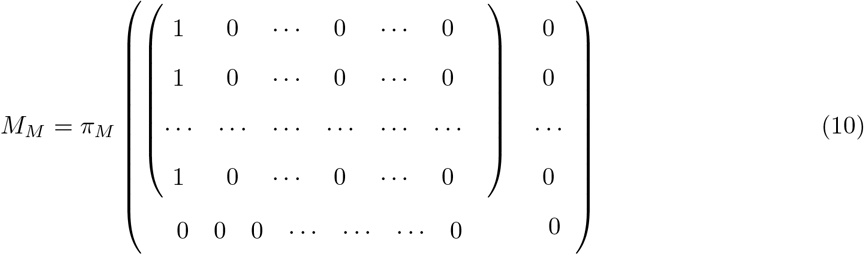

And third, the interphase matrix *M_I_* exploits the asynchronous dataset to infer the average expression levels during interphase. The matrix takes the following form:

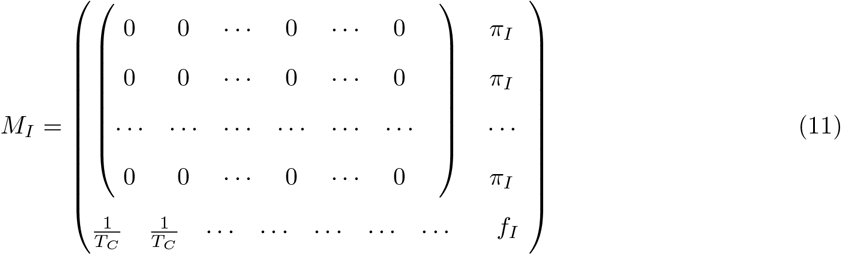

where the cell cycle duration *T_C_* is set to 24*h* [22].

Hence, the deconvolution problem can be understood as a multiple linear regression and, therefore, we can infer the gene expression in the space of the cell-cycle pseduotime by optimizing the following quadratic loss function:

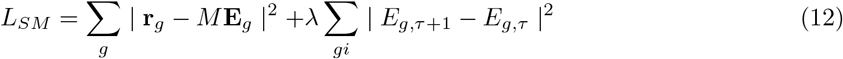

where we added a smooth Ridge regularization term to be able to solve the overrepresented linear model and avoid overfitting. Then, the solution is 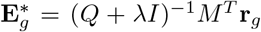, where *Q* = *M^T^ M* and *I* is the regularization matrix.

Finally, To choose the parameter *λ*, we calculated the Akaike Information Criterion (AIC) and the Bayesian Information Criterion (BIC) scores in function of *λ*, as follows [23, 24]:

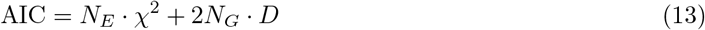

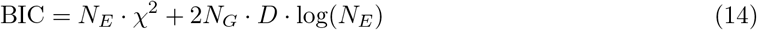

where *N_E_* is the number of experimental time points, *N_G_* the total number of genes, 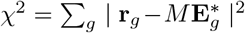 is the minimum error and *D* is the degree of freedoms that, for a multiple linear regression model with smooth Ridge regularitation, can be calculated as *D* = Tr(*M* ((*Q* + *λI*)^−1^*M^T^*). The BIC score tents to introduce a stronger penalty producing a solution more robust against overfitting therefore we chose *λ* = 0.79 that minimizes the BIC score (see Fig. S1).

### Visualization of the gene expression through heatmaps

To represent the gene expression as shown in Fig. 1 panel A, processed EU-RNA-Seq data at the transcript level from [10] were used. Transcript FPKMs from the same gene were then grouped to obtain gene level EU-RNA-Seq data, and all the genes with a low expression on the asynchronous sample (< 36 FPKM) were excluded (see Fig. S1, panel A). Then, a Z-score was calculated, correcting each FPKM value by subtracting the mean *μ_g_* and dividing by the standard deviation *σ_g_*, both *μ_g_* and *σ_g_* calculated over the corresponding gene. Genes were divided into 5 clusters (the optimum number to obtain significantly different profiles) according to their Z-score over time, by using the *KMeans* tool from *sklearn* python library. The heatmap was represented by using *seaborn* python library, ordering the genes of each cluster according to their norm with respect to the corresponding cluster average expression.

### Inference of transcription factor activities

We developed an ISMARA-like model [13] where the expression of a given gene with respect to the cell-cycle pseudotime can be obtained as a linear combination of time-dependent activities of all TFs that can potentially bind its promoter. First, a as proposed in [13], we preprocessed our data as follows: first, to revert the z-score transformation performed above we multiplied the gene expression values *E_gα_* by the standard deviation *σ_g_* and added the average *μ_g_*. Second, in order to calculate the log2 expression for all genes, we add a pseudo-count to the corrected *E_gτ_* values, i.e. for every given time *τ*, we ranked all the values higher than zero and we calculated the 5th percentile *pc_τ_*. We then added *pc_τ_* to the corresponding *E_gτ_*. After that, we calculated 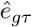, i.e. normalized values of the gene expression at a cell-cycle pseudotime *τ*, as follows:

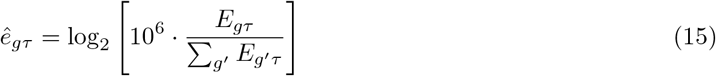

we further normalized the expression of genes across pseudotime and genes resulting in 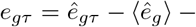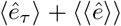. Finally, we write the linear model as:

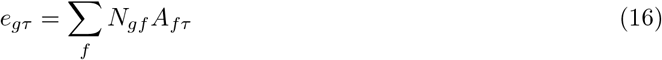

where the value *N_gf_* represents the number of binding sites for the TF *f* on the gene promoter *g*, taking into account the affinity between the motif of *f* and the sequence of the promoter; and, the unknown parameter *A_fτ_* is the activity of the TF *f* at a given cell-cycle pseudotime *τ*. The binding site matrix is further normalized to ensure ∑_*g*_ *N_gf_* = 0. Note that the TF activities are then zero mean variables.

Then we used least square fitting to obtain the TF activites. To avoid overfitting we included a Ridge regularization penalty. To estimate the weight of the regularization we calculated the Mean Square Error (MSE) for a training and a test datasets and we performed a 80-20 cross-validation. A regularization factor *λ* = 443 was chosen, corresponding to the minimum of the MSE of the test dataset (see Fig. S2). In addition, we calculated the explained variance (EV) of the model, 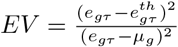, where 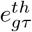 is the theoretical expression of the gene *g* at internal cell cycle time *τ*, i.e. calculated using the inferred activity *A_fτ_* and the matrix *N_gf_*, and *μ_g_* is the mean among all the values *e_gτ_*. We obtained a regularization factor *λ* = 443, corresponding to the maximum of the EV of the test dataset Fig. S2, in accordance with the minimum obtained for the MSE.

### Visualization of the TFs activities through heatmaps

To represent the TFs activities as shown in Fig.2, the TFs were divided into 3 clusters according to their activity dynamics over *τ*. First, we calculated the standard deviation over time of TF activities as 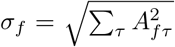 and classify a TF as high amplitude dynamic if *σ_f_* > 0.07. Second, we sorted TFs according to when their maximum activity peak occurred and defined a TF as mitotic active if the peak appeared before *τ*_mit_. Therefore, TFs were classified as either mitotic active, early-G1 active or non-dynamic. The heatmap in Fig.2 was represented by using *seaborn* python library, ordering the TFs in each cluster by the the first reached maximum over *τ*. To represent the TFs activity as shown in Fig. S3 only TFs corresponding to genes belonging to the Gene Ontology (GO) category (Cell Cycle - GO:0007049) were considered.

### Core Regulatory Network

To build the core regulatory network (CRN) we selected the TFs that showed a high degree of explanatory power to reproduce the gene expression dynamics. To do that, we assigned a score for each TF based on its contribution to the explained variance by calculating a *reduced* explained variance 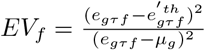, i.e. the EV as shown in the section Inference of transcription factor activities but removing from the model the corresponding TF *f*. Then, we defined the importance score of a TF by the ratio 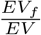 and we ranked all TFs according to this score taking into account that the smaller is the ratio the higher is the impact of the TF on the explanatory power of the model. The figure was then generated by using digraph library in Matlab, by selecting relevant TFs and corresponding genes in the N matrix.

### Genes expression dynamics and bookmarking

To establish which genes are associated to FOXA1, as shown in Fig. 3, panel A, we took into account the mitotic ChIP-Seq peaks of FOXA1 from [6]. Then, for every peak, we selected the nearest expressed gene, using as references the corresponding TSS and the average point of the selected peak. So, we obtained a list of expressed genes that we defined the genes bound by FOXA1 during mitosis.

To establish which genes tend to be regulated by TFs with high or low MBF [4], as shown in Fig. 3, panels C and D, we calculated what we called *MBF weighted average scores* (MWAS) as follows: for each gene *g*, we took the number of binding sites *N_gf_* corresponding to the factors *f* for which we know the MBF score. Each of these values was multiplied for the corresponding MBF, and then they were summed all together. Finally, this sum was divided by the total number of binding sites, i.e. the sum ∑_*f*_ *N_gf_*. This score is what we called MWAS. Then we ranked the genes according to the MWAS, and we removed the ones with MWAS = 0. The 10% of the genes with the highest MWAS were then considered associated with enriched TFs (”enriched genes”), while the 10% of the genes with the lowest MWAS were considered associated with the depleted TFs (”depleted genes”).

To obtain genes enriched or depleted in binding sites, we calculated for each gene promoter the total number of binding sites, i.e. ∑_*f*_ *N_gf_* and then the 10% of the genes with largest number of binding sites and the 10% of the genes with smallest number were considered to calculate average expression profiles as shown in Fig. 3 panel E and F.

## Supporting information

**Fig. S1.**
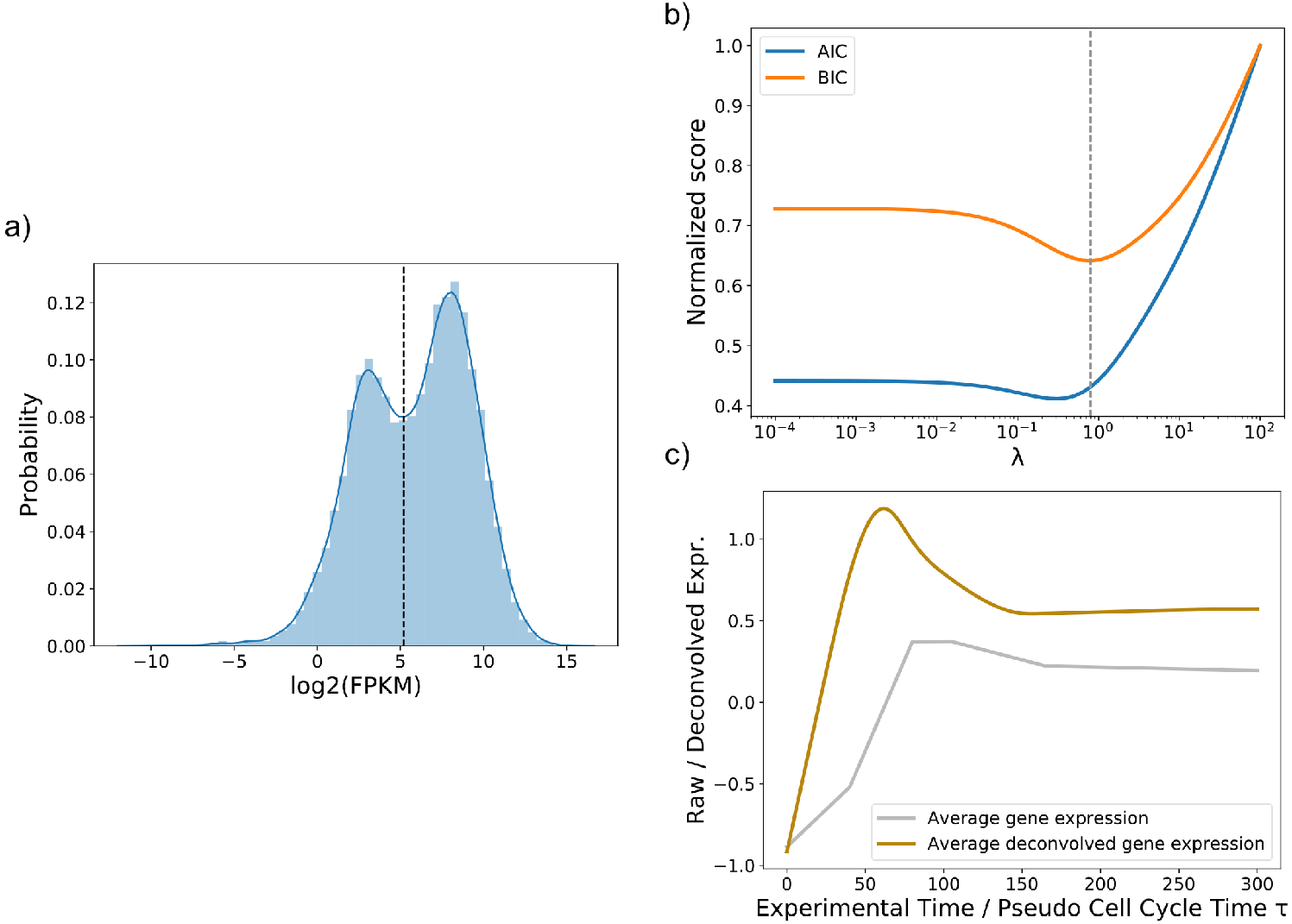
Data processing for deconvolution. **A**: Log2 histogram of FPKM reads at gene level for asynchronous data. The dashed vertical line represents the threshold we considered to process our data: genes with asynchronous *FPKM* < 36.76 were excluded. **B**: AIC and BIC scores were calculated in order to establish the best *λ* parameter for the regularization of the deconvolution process (see Methods). Both AIC and BIC showed a minimum, and we choose *λ* = 0.79, corresponding to the BIC minimum (dashed vertical line). **C**: Average gene expression of convolved (grey line) and deconvolved (yellowish line) data were represented on the same plot.

**Fig. S2.**
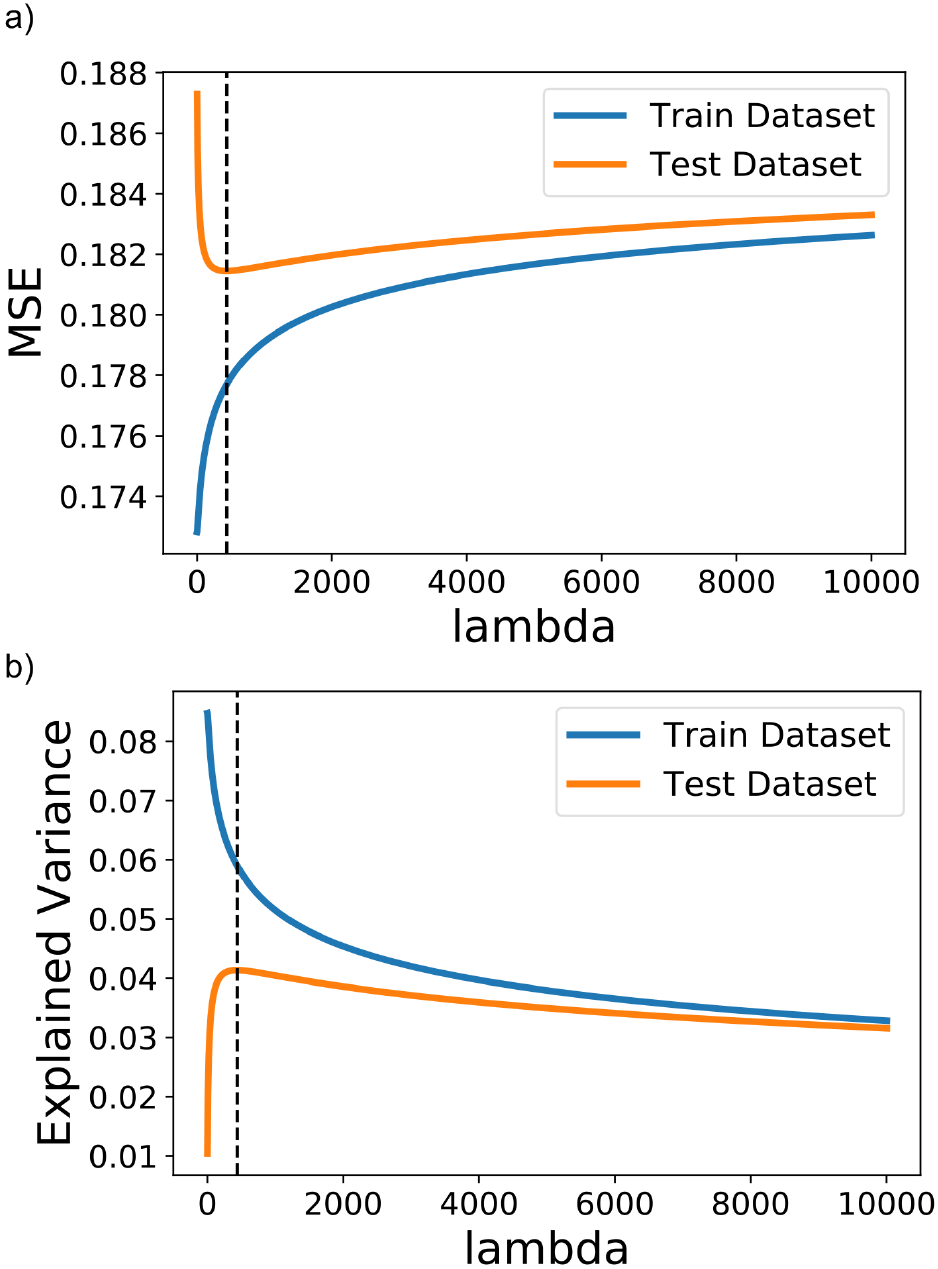
Cross validation of the linear model. **A**: A cross-validation 80/20 was performed to find the best *λ* regularization parameter for inferring the TFs activity (see Inference of transcription factor activities). A value *λ* = 443 was chosen (dashed vertical line), corresponding to the minimum of the Mean Squared Error (MSE) of the test dataset (see Methods). **B**: The same analysis shown in the panel A was performed by using Explained Variance (EV) instead of MSE. The dashed vertical line corresponds to the maximum *λ* = 443, in accordance with the minimum MSE.

**Fig. S3.**
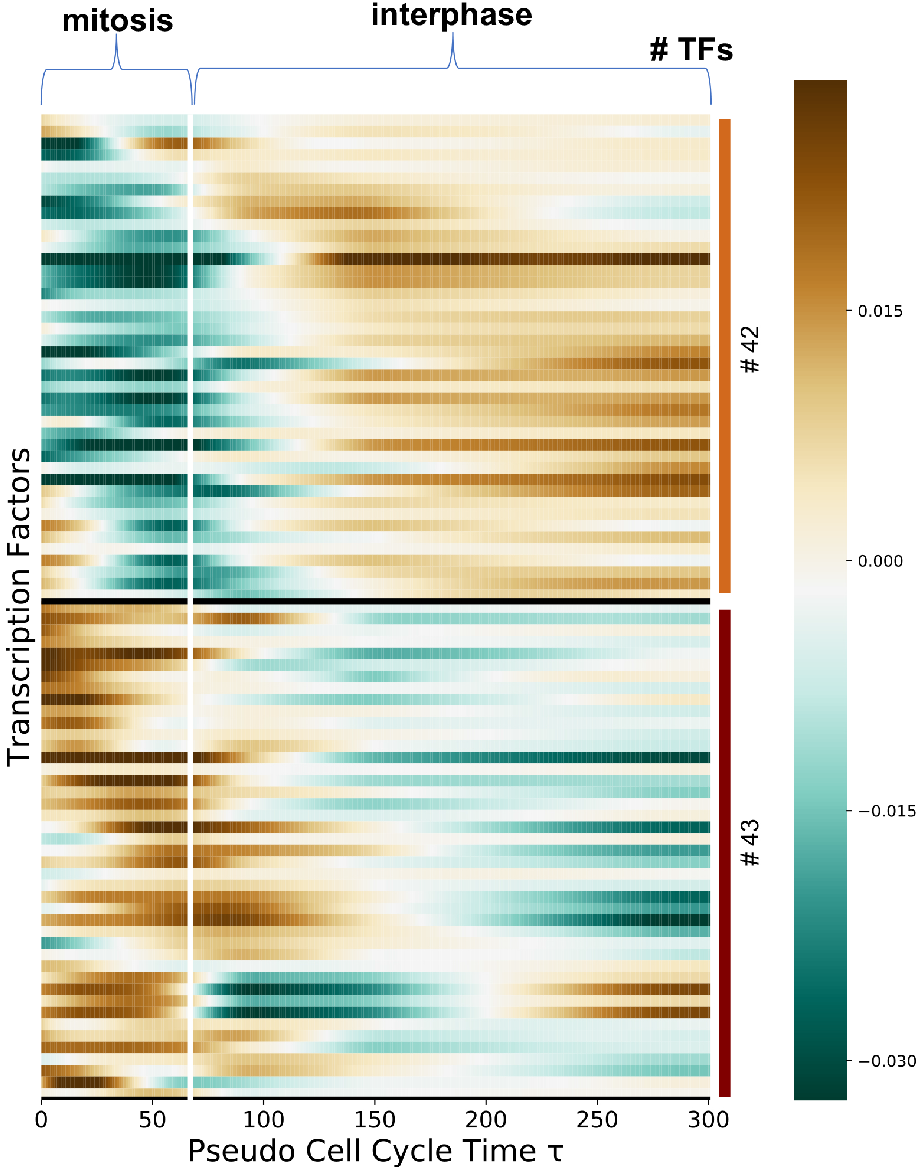
Transcription factors dynamics taking into account only cell-cycle GO category. Here, only TFs associated to genes belonging to the Gene Ontology (GO:0007049) category have been shown and clustered. In this case, only 2 main groups of TFs have been individuated, and both of them show a significant activity change over *τ*. The vertical white line represents *τ*_mit_, and separates ideally the mitosis from the interphase. On the right, the number of TFs for every cluster is indicated.

**Fig. S4.**
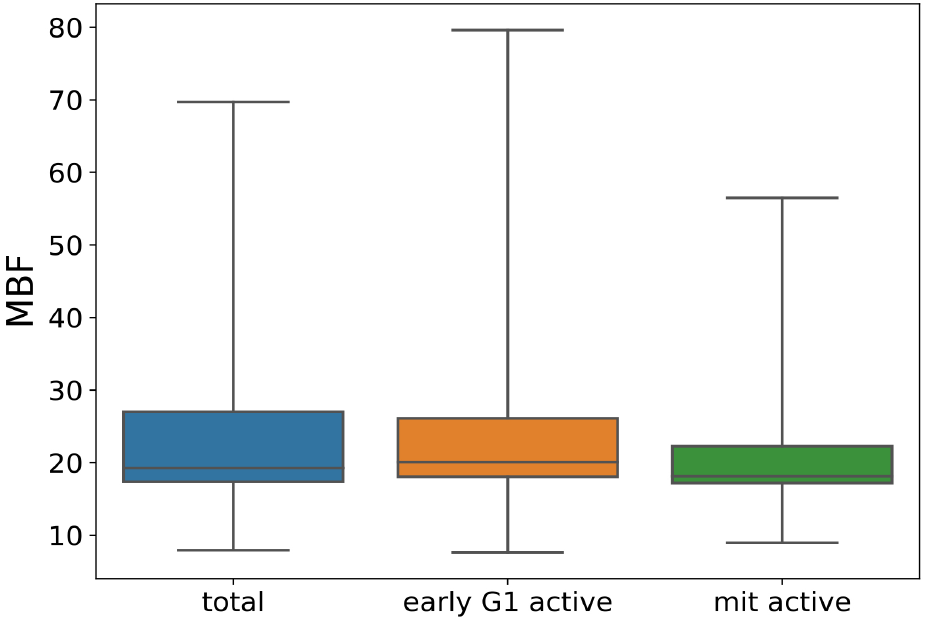
Average MBF for mitotic and early G1 active transcription factors. Boxplots showing the average MBF for TFs with higher activity during mitotis (green box) and during early G1 (orange box) respectively, in comparison with the average of all TFs (blue box).

## Acknowledgments

This work was possible thanks to the funding from the LabEx INRT grant and the IDEX actrativité program of the University of Strasbourg. Furthermore, S.S., A.R. and N.M are grateful to the IGBMC to provide an excellent working environment.

